# DevoWorm: differentiation waves and computation in C. elegans embryogenesis

**DOI:** 10.1101/009993

**Authors:** Bradly Alicea, Steven McGrew, Richard Gordon, Stephen Larson, Tim Warrington, Mark Watts

**Affiliations:** Orthogonal Research, Champaign, IL USA; New Light Industries, Ltd, Spokane, WA USA; Embryogenesis Center, Gulf Specimen Marine Laboratory, Panacea, FL USA; C.S. Mott Center for Human Growth and Development, Department of Obstetrics and Gynecology, Wayne State University, Detroit, MI USA; MetaCell, LLC, San Diego, CA USA; Department of Molecular Biology and Biochemistry, Simon Fraser University, Burnaby, BC Canada; Department of Computer Science, University of Texas at Austin, Austin, TX USA

## Abstract

Development is a complex process that, under normal circumstances, proceeds in a stable and patterned fashion. Developmental morphogenesis (called embryogenesis) can tell us a great deal about the function and structure of an adult organism. One of the most important aspects of development to understand is the progression of cell division and differentiation in what will become an adult worm. This is where the DevoWorm project can both address multiple outstanding theoretical issues and provide graphical clarity to the embryogenetic process. As a representative of mosaic development, *C. elegans* embryogenesis is both tractable in terms of cell number and relatively well-characterized. In this paper, we will lay out a theoretical re-interpretation of embryogenesis in addition to developing an RDF-based computational framework for visualizing the results of this theoretical effort. Our theoretical efforts will ultimately involve the construction of a differentiation tree and data analyses that support the concept of differentiation waves acting to coordinate cellular differentiation and embryonic form. The differentiation tree will also feature a means to perturb development in a manner that mimics phenotypic mutagenesis. This will allow us to understand the selective variability that is inherent in biological development, but that remains so poorly understood. In tandem, these developments will allow us to construct a conceptual and computational framework which can be applied to both mosaic and regulative development.

## Introduction

In many respects, *C. elegans* is the perfect model organism for understanding the process of cellular differentiation in development. The *C. elegans* embryo develops from a single undifferentiated cell to 959 cells in the adult hermaphrodite [1] and 1031 cells in the adult male [2]. While the amount of developmental complexity is tractable from an analytical standpoint, two other features of *C. elegans* development are ripe for revisitation. While the number of cells generated in development varies by academic source and the worm’s sex, the number of cells that are generated during embryogenesis, undergo apoptosis, and are functionally unique is both relatively small and well-characterized. For example, Sulston [3] report that 671 cells are generated in embryogenesis, 111 of the cells in the male and 113 in hermaphrodites undergo apoptosis, and approximately 560 are involved in producing terminal adult cells. Other counts [1] report 959 cells for an adult hermaphrodite, with around 850 unique cells and around 50 cells as equivalent pairs.

Furthermore, each cell plays a specific functional role which does not change in the adult form or from individual to individual. This allows for a rather detailed accounting of developmental symmetry along three anatomical axes. In *C.elegans*, cell lineages unfold pairwise from founder cell to organ along one of three axes. This allows us to track individual cells and lineages throughout development, in addition to attaching relational data to each cell. Developmental invariance also allows for us to build a strictly hierarchical tree based on cell-specific differentiation events across the entire body and developmental process [4].

While this is an attractive model, it may not fully account for the developmental process. For example, rather than unfolding in a linear fashion, embryogenesis is marked with significant false starts (apotosis) and creative destruction (autophagy). While the work of Sulston, Albertson, and Thomson [4] provides a means for differentiation, the processes that underlie differentiation patterns remain unaccounted for. As their lineage tree is organized from head to tail, the tree structure that results does not account for the dynamics of embryogenesis. Reorganizing the lineage tree into a differentiation tree may allow us to triangulate the exact meaning and nature of what it means to differentiate. Furthermore, detailed spatial relations, such as the 2- and 4-fold symmetry mentioned earlier, can provide an accurate emulation of developmental processes and organismal-scale biology. This will provide a means for OpenWorm users and *C.elegans* biologists alike to better understand the “how” and “why” of *C.elegans* developmental biology.

We propose that the differentiation tree (Figure 1) can be used to re-interpret the lineage tree model of mosaic development [5]. Differentiation trees provide a model that attempts to explain both mosaic and regulative development. Examples of regulative and mosaic development are found in nematodes and amphibians, respectively. In regulative development, an embryonic tissue is divided into two new embryonic tissues by a pair of waves. A schematic of the differentiation wave concept is shown in Figure 2. A contraction wave traverses one portion of the tissue while an expansion wave traverses the remaining part. The analogues of this in mosaic embryos, in which differentiation occurs via asymmetric cell divisions, are shown in Table 1.

**Table 1.** Differences between regulative and mosaic embryonic development.

| **Regulative** | **Mosaic** |
| --- | --- |
| Contraction wave | Small cell in an asymmetric division |
| Expansion wave | Large cell in an asymmetric division |
| Cell proliferation without further differentiation | Symmetric cell division yielding two equal sized cells of the same type as their mother cell |

**Figure 1.**
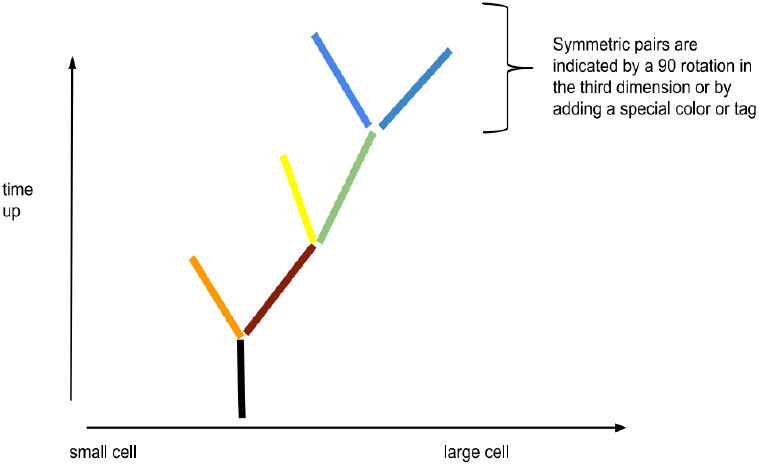
An example of how the differentiation tree might be applied to *C. elegans* development.

**Figure 2.**
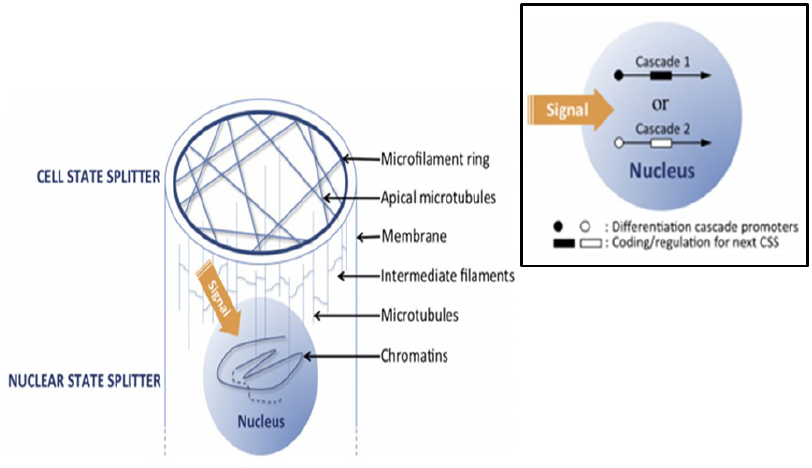
An example of a differentiation wave’s mode of action on a single cell. **MAIN:** cellular and cytoskeletal sites of action, **INSET:** nuclear sites of action. COURTESY: Figure 1 in [7].

One goal of this project is to reorder cells in the lineage tree from tail/head ordering to small/large ordering to create the differentiation tree of *C. elegans*. This will be done to make the tree consistent with the mosaic type of development. The strength of the differentiation tree approach is demonstrated in Proposition 170 of [5]. Namely, combining the differentiation tree with mosaic development allows us to infer the cellular state and angle of differentiation for any cell in the differentiation tree.

## Approach

Our approach is intended to be twofold: a theoretical re-interpretation of *C. elegans* embryogenesis, and a computational presentation of embryogenetic structure. The first aim of our project is to re-interpret *C. elegans* embryogenesis and the lineage tree framework as a process that can be summarized in the form of differentiation trees and differentiation waves. This will involve analysis of heteogeneous data, including semantic, microscopy, and gene expression data. The second aim of our project will be to capture the bifurcation process through a computational architecture known as the Resource Description Framework (RDF). This will allow us to build a descriptive network using a common language of tags, which can be use to label cells with metadata in an extensible manner. This produces graph structured models of the data mapped onto the differentiation tree structure. The dual nature of the project is a strength in that the theoretical framework and analysis will inform the informatics component, and vice versa.

The first part of our approach is to establish a computational roadmap to testing and developing theoretical propositions. Specifically, we will focus on identifying the correlates of differentiation trees and waves in the developmental (and adult) *C. elegans* data. The application of differentiation waves to *C. elegans* development is further discussed in [6], while details about the propagation mechanisms behind differentiation waves can be found in [7].

### Expansion/Contraction Wave Sites of Action

In general, differentiation trees are based on the presence of expansion and contraction waves triggered by cell state splitter activity in individual cells. A cell state splitter: cytoskeletal structure hypothesized to send a binary signal. This is important change of state information that is transmitted to genomic regulatory mechanisms. This should allow a cell to transform into one of two new cell types. In this way, the cell state splitter triggers a differentiation signal which can be transmitted form cell to cell in a population or tissue.

### Broader Implications

Mechanical signals are not the only possible mechanism for the cell state splitter. In C. elegans (mosaic development) for example, the candidate mechanism could be mechanical resonance, juxtacrine signaling, cell motility, or a combination of these factors. In this way, differentiation trees and differentiation waves are an extension of reaction-diffusion morphogenetic models advanced by Turing [8]. However, Turing’s model scales poorly when executed amongst large cell populations [9], becoming computationally prohibitive. As they are consistent with more biologically realistic models of morphogenesis [10], discovering the empirical correlates of differentiation trees and waves is a significant advance in the study of embryogenesis.

The second part of our approach is to use computational tools to uncover the structure and context of *C. elegans* embryogenesis. These include specialized RDF data structures and XML graphical structures. As a layered rendering of heterogeneous data supporting the differentiation tree/waves concept, these data structures will provide an interactive and extensible framework for generalization and future work.

### N-Quad Data Structures

To better represent information about worm development, we can use an RDF data structure called an N-Quad (Table 1). N-Quads are simple statements that consist of a 4-tuple (a subject, predicate, an object, and context). Simpler N-triples can be used to represent standard 3-tuples of information. For example, we can use a 3-tuple (*x*,*y*,*z*) to describe the initial location of a particular *C. elegans* cell in space. Another 3-tuple consists of three parameters most fundamental to this version of the project: cell size (*i*), division event number (*t*), and the spatial angle of differentiation (two components – *θ, φ*). By using this framework, representing functional attributes such as gene expression values and specific physiological attributes will also be possible as the data become available [10].

The first database we will use is information about what happens in development and the position of each cell with respect to the developmental process. To map this information from raw data to the differentiation tree, we need representations that allow us to determine the order of branching and construct a hierarchical differentiation tree structure. We can then conduct a more focused inquiry of *C. elegans* development such as determining formal mechanisms behind the asymmetric cell divisions. This framework also allows us to correlate developmental events amongst physical neighbors in the adult phenotype. We can also use the RDF-based annotations to postulate candidate developmental mechanisms amongst different cells in the model. As lineage trees are altered in some *C. elegens* mutants, the corresponding mutant differentiation trees may provide insights into the morphogenetic roles of mutants.

### Graph Structure

The Wormbase dataset [11] provides us with a multi-dimensional topology within which to place each cell, providing an underlying structure to the spatial semantic graph in Table 2. This may also allow us to determine whether or not cells migrate together, and exactly how short and long range order gets maintained during development. Yet we are still dealing with semi-structured data. Thus, our data and its annotations will be integrated and presented (e.g. visualized) using the Unified Data Access (UDA) Layer protocol written in PyMol. The leading candidate for creating these augmented network structures is NetworkX [12]. This will allow for additional relational attributes to be extracted and presented in a highly visual format.

**Table 2.**
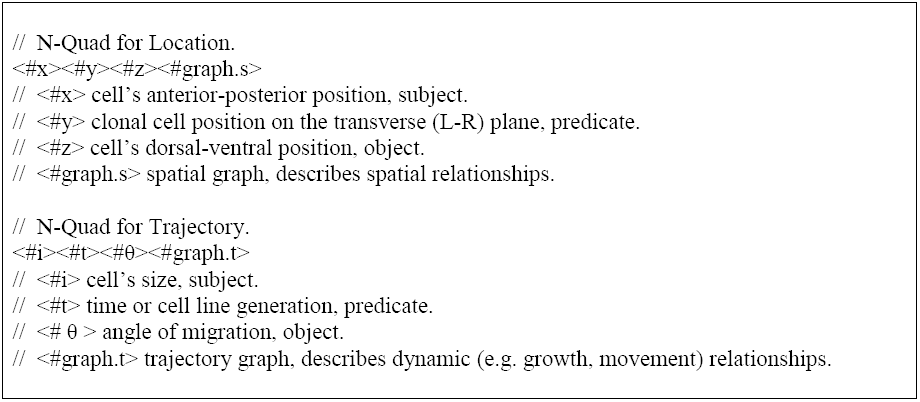
Two examples of the N-quad data structure for a single, hypothetical cell. Each <string> attribute consists of a parameter value.

### Classes, Methods, and Additional Features

We will be using various existing and customized classes and subclasses to characterize *C. elegans* development. In some cases, these classes can come directly from the OpenWorm platform. In other cases, custom object types will have to be created. One example of this is the “neuron” object, which is a class in OpenWorm but may become a subclass in DevoWorm (see Table 3). We also want to characterize 3-dimensional position relative to the founder cells and changes in volume after a pair of daughter cell is created from their mother cell. Additional functionality such as cell annotations or links to relevant literature may also be possible.

**Table 3.**
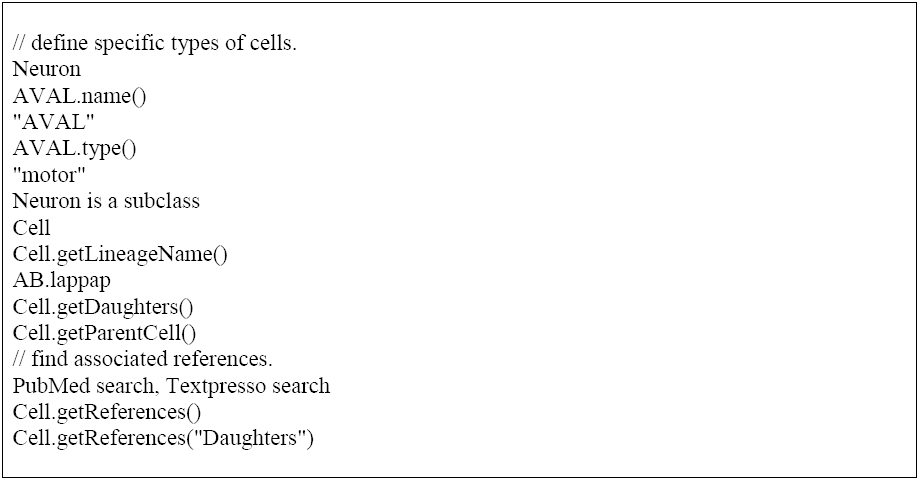
Pseudo-code for defining specific cells as objects in DevoWorm.

### Data Sources

Our relational and semantic data [13] come from the OpenSource project database, which in turn comes from the work of [3] and [4]. In the original dataset, a convention is used that allows for the lineage of individual cells to be identified (e.g. ABpldaar). This naming convention is highlighted in Table 4. The identity of founder cells (analogous to blastomeres in vertebrates) is determined by capitalized prefixes (e.g. AB) that define each lineage. Cells within this lineage are further specified by a lower-case suffix (e.g. pldaar), the letters representing the sequential orientation and direction of division. These data will be co-registered with micrograph data, courtesy of the White Lab and Sharon (Fong-Mei) Lu and the University of Wisconsin. These data include 12 focal planes of 2 embryos from the male-feamle gonadal stage to the 100-cell stage. This should provide us with even more precise positional and temporal information.

**Table 4.**
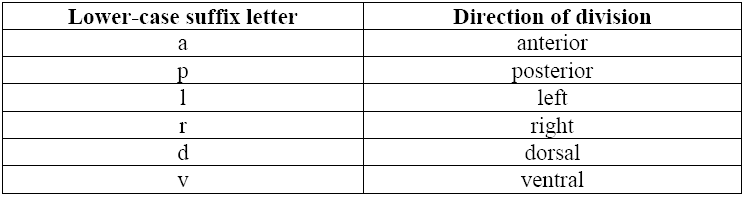
Convention for cell names [3] extracted from the lineage tree of Sulston, Albertson, and Thomson [4].

| Lower-case suffix letter | Direction of division |
| --- | --- |
| a | anterior |
| p | posterior |
| l | left |
| r | right |
| d | dorsal |
| v | ventral |

Our gene expression dataset is secondary data from [14, 15]. The first dataset [14] contains the expression of selected protein coding genes for different timepoints in development. The source of these data is the Gene Expression Omnibus database (http://www.ncbi.nlm.nih.gov/geo/), Accession Number GSE2180. The second dataset [15] contains both gene- and cell-specific expression data during development. The source of these data is the Expression Patterns in *C. elegans* (EPIC) database (http://epic.gs.washington.edu/). Taken together, these data will serve to demonstrate various attributes of developmental processes that are potentially related to differentiation trees and waves.

## Analysis of C. elegans Embryogenesis

### Shape Analysis and Developmental Trajectory

An alternate way to understand lineage tree data as the products of differentiation waves is to analyze the shape of cellular differentiation in a metric space. In this case, we visualized the differentiation of cells relative to lineage founder cells in lineage space. Lineage space is a low-dimensional representation of the *C. elegans* embryo at an advanced stage of embryogenesis, and is centered the founder cell of one (or a composite of all) lineages in the worm. Calculating a cell’s position in lineage space involved the following: the net cell position for each datapoint was obtained and divided into the maximum distance along the axis. This maximum distance is dataset-dependent, and creates a metric space where 0.0 is the founder cell location, and all cells found in the embryo can be found along interval {1.0, -1.0} for the A-P, L-R, and D-V axes. Both the angles and Euclidean distances were calculated relative to the founder cell. These variables give us an abstract approximation of how development unfolds in space for a single lineage(AB).

We can further analyze these data by visualizing the lineage space using angle of differentiation and Euclidean distance from the founder cell. These variables were then visualized for the AB lineage using a polar plot (Figure 3). Due to the lack of variability along the D-V axis, only the L-R and A-P axes were used. Nevertheless, Figure 3 demonstrates the asymmetries of this lineage space for all lineages in a *C. elegans* embryo using lineage tree data from [3]. This graph also shows the spread of cells relative to the maximal number of cell divisions in the embryo. In the AB lineage, there is skew in the anterior direction, particularly along the midline (relative to the founder cell position). This analysis also demonstrates a dispersal of the lineage in space, an outcome that might be predicted by the propagation of a differentiation wave.

**Figure 3.**
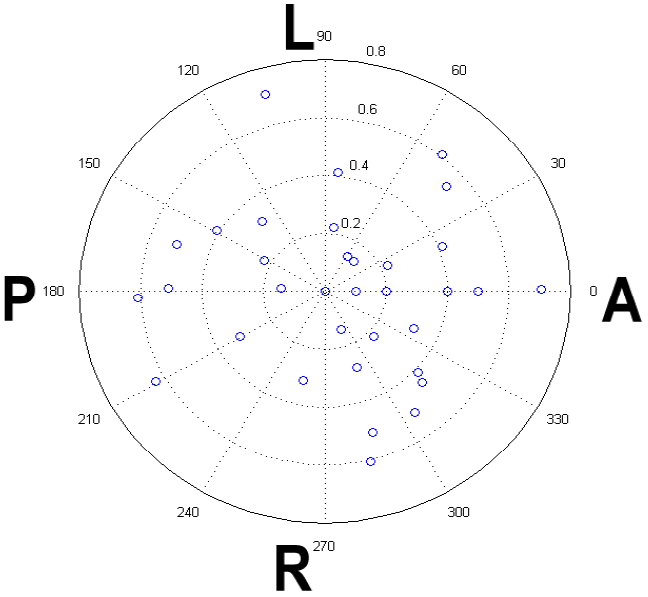
Polar plot of cell locations in lineage space using developmental dataset. The datapoints represent a cell’s angular orientation relative to the founder cell position in lineage space and its Euclidean distance from the founder cell position in lineage space.

### Gene Expression and Developmental Age

We can also estimate the developmental age of cells in an adult phenotype through measuring peak gene expression activity. We used a dataset that consisted of mature cells of all lineages found in the adult phenotype, and consisted of 671 distinct cells, 5 lineages and 18-73 genes per cell. The authors of [15] define peak expression as the maximal reporter intensity observed in cell of measurement or any of its ancestors. This has the effect of carrying forward expression patterns to terminal set of cells in a lineage, resulting in a cumulative effect of residual reporter protein from both ancestral cells and endogenous expression. While endogenous expression predominates, the overall intensity differences between cells denote both age and symmetrical cell division.

From the graph in Figure 4, we can see that there are subtle patterns of variation with respect to spatial-dependent gene expression patterns. While the first principle component reveals some variance structure with respect to a cell’s location in space, there is no clear relationship between lineage age or lineage identifier and a cell’s position in lineage space. Future directions might involve a whole-genome analysis at the single cell level, which would allow us to find genes that are strongly associated with a cell’s location in physical space and the lineage hierarchy.

**Figure 4.**
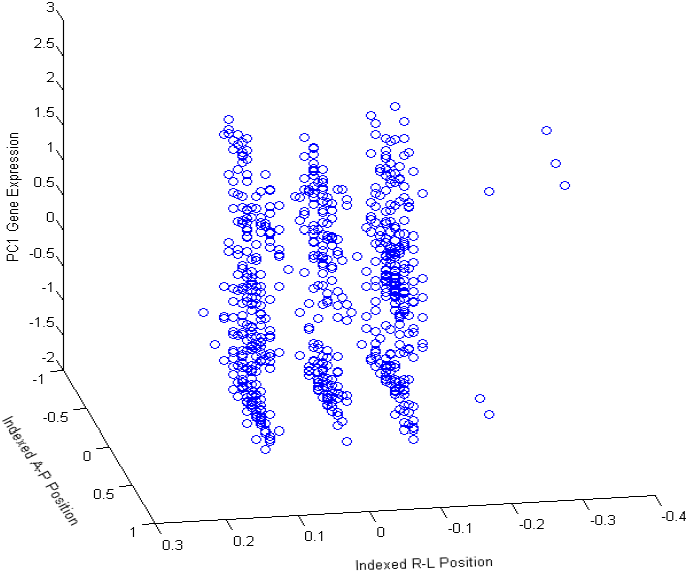
Three-dimensional plot of variability in lineage space along the L-R and A-P axes against PC1 for gene expression.

## Conclusion

As a conceptual and technological initiative, the DevoWorm project will ultimately provide a window into the complexity and causal mechanisms of development. Our focus on theoretical re-interpretation will allow us to bring theoretical concepts to a more general audience. Although there is great potential to learn more about the fundamental principles of cellular differentiation and patterned development, one potential application of this work is educational outreach. We also aim to construct an extensible visualization framework for not only representing data on *C. elegans* development, but also for advancing the theory behind incompletely-observed developmental dynamics in Eutelic animals [16]. We plan to do this through the use of heterogeneous data, computational modeling, and the application of data structures. Additionally, we wish to address meta-issues such as the role of variability and effects of perturbation on the developmental process. In addition to the overarching goal of emulating the developmental biology of an entire organism with a minimal amount of information [17], it is hoped that the DevoWorm project provides both theoretical and empirical resolution to profound biological problems.

